# The maximum solubility product marks the threshold for condensation of multivalent biomolecules

**DOI:** 10.1101/2022.10.04.510809

**Authors:** Aniruddha Chattaraj, Leslie M. Loew

## Abstract

Clustering of weakly interacting multivalent biomolecules underlies the formation of membraneless compartments known as condensates. As opposed to single component (homotypic) systems, the concentration dependence of multi-component (heterotypic) condensate formation is not well understood. We previously proposed the solubility product (SP), the product of monomer concentrations in the dilute phase, as a tool for understanding the concentration dependence of multi-component systems. In the current study, we further explore the limits of the SP concept using spatial Langevin dynamics and rule-based stochastic simulations. We show, for a variety of idealized molecular structures, how the maximum SP coincides with the onset of the phase transition, i.e., the formation of large clusters. We reveal the importance of intra-cluster binding in steering the free and cluster phase molecular distributions. We also show how structural features of biomolecules shape the solubility product profiles. The interplay of flexibility, length and steric hindrance of linker regions controls the phase transition threshold. Remarkably, when solubility products are normalized to non-dimensional variables and plotted against the concentration scaled to the threshold for phase transition, the curves all coincide independent of the structural features of the binding partners. Similar coincidence is observed for the normalized clustering vs. concentration plots. Overall, the principles derived from these systematic models will help guide and interpret in vitro and in vivo experiments on the biophysics of biomolecular condensates.

**Significance Statement:** Biomolecular condensates are macroscopic intracellular structures that are composed of weakly interacting macromolecules. Because their composition can be complex, there are no simple rules for how condensates form as a function of the concentrations of their individual components. In this work, we show how the solubility product (SP), the product of monomer concentrations in the dilute phase, might serve as a tool for predicting the concentration dependence for condensation of multi-component systems. Specifically, Langevin dynamics simulations of the clustering of a series of multivalent binding partners reveals how the maximum SP is always attained at the same concentration as the appearance of large clusters. Experimental application of the SP concept should help rationalize the cellular formation of biomolecular condensates.

## Introduction

Biomolecular condensates are membrane-less sub-cellular compartments that play an important role in spatiotemporal regulation of cellular biochemistry [1]. Clustering of weakly interacting multivalent biomolecules (proteins and nucleic acids) leads to condensate formation via phase separation [2-5]. Condensate biology has developed explosively, at least in part because of its potential connections to a wide variety of diseases [6-8].

Biomolecular phase separation combines two distinct albeit coupled physical processes: density transition and network transition [9]. A homogeneous polymer solution, above a threshold concentration, splits into a polymer-rich dense phase (condensates) surrounded by a solvent-rich dilute phase. This density transition is generally explained by the classical Flory – Huggins theory [10] where a solubility parameter quantifies the competitive interactions between the polymer and solvent. Multivalent biomolecules also engage in biophysical interactions with other molecules via specific binding motifs in their sequences. These multivalent interactions underlie molecular network (cluster) formation and the transition from a dispersed phase to a clustered phase is referred to as gelation or percolation transition [11, 12]. Since clustering of biomolecules reduces the solubility of the system, the percolation transition is generally strongly coupled with the density transition. In this paper, we use the molecular clustering process as a useful proxy to predict the phase separation tendency of the system. We use the term “phase transition” to denote the jump between the dispersed phase (monomers and small oligomers) and the clustered phase (monomers, small oligomers and very large clusters).

For a single component (homotypic) system, the dilute phase concentration of the biomolecule remains constant beyond the concentration threshold while the condensates grow in size and numbers [13]. Similarly, for a multi-component heterotypic system, the solubility product (SP, the stoichiometry-adjusted product of the dilute phase monomer concentrations) converges to an approximately constant level when the system starts to form large clusters, as we established in previous work [14]. In other words, as the total concentration of the molecular components increases, the phase transition boundary of the system is marked by a maximum SP that remains approximately constant, the solubility product constant (Ksp). In our previous study [14], we primarily used a non-spatial stochastic network-free simulator, NFsim [15], to demonstrate the statistical effects of valency on the phase transition threshold. In this paper, we study the effect of molecular structural features on the solubility product profiles, using the SpringSaLaD spatial simulator [16] and explore the limits of the Ksp concept that emerged from our previous non-spatial simulations. We introduce a pair of non-dimensional parameters (normalized SP and normalized ACO) that can serve as dimensionless metric to compare these multivalent systems undergoing phase transition. We further present new statistical analysis to better quantify the binding saturation and compaction of multivalent clusters.

## Materials and Methods

### Spatial Langevin Dynamics

We use the SpringSaLaD simulation platform [16] to probe the effects of molecular structures on the solubility product profiles. Coarse grained molecular dynamics has been employed by multiple previous studies [17-21] to investigate molecular clustering in the context of phase transition. In this framework, a macromolecule is modelled as a “bead-spring” polymer where certain beads engage in a binding interaction (‘stickers”) with their cognate sites while other beads serve as flexible tethers (“spacers”). SpringSaLaD models biomolecule in a similar fashion where a biopolymer is a collection of “binding” and “linker” sites. Although the algorithm and its application has been previously described [14, 22], we will briefly summarize the workflow here. Each biomolecule is modeled as a collection of hard spheres connected by stiff spring-like linkers. Particle motion is governed by the overdamped Langevin equation where a site experiences two types of forces – a resultant of harmonic forces transmitted by the bonded neighboring sites and a stochastic force calibrated with the input diffusion coefficient to produce Bronwnian motion. A binding site within a molecule reacts with its partner binding site by a set of users defined rules. For individual pairwise binding, users provide a macroscopic on rate and off rate based on experimental observations. Using the on-rates, the software computes microscopic binding probabilities, applied once the binding spheres come within a computed reaction radius, based on the user-defined rate constants, diffusion constants and physical radii of the binding site-pairs. Therefore, if the interacting spheres are already within a cluster due to multivalent binding, their mutual translational freedom is lost and they will be more likely to find themselves within the reaction radius. Excluded volume is also enforced by individual sites.

We construct the model using the graphical interface of the software and then run multiple simulations in parallel using the High Performance Computing facility at UConn Health (https://health.uconn.edu/high-performance-computing/). A typical 50 ms simulation (time step = 10 ns) of a system consisting of 6000 sites (400 molecules) takes about 24 hours on a Xeon processor. The execution time, of course, is very sensitive to the simulation conditions like number of molecules, structural constraints within molecules etc.

### Non-spatial simulations

We use the rule based stochastic simulator, NFsim [15] to isolate the statistical effects of valency on molecular clustering. As described earlier [14], NFsim represents a molecular object as a collection of binding sites. A set of user defined rules specify the binding kinetics between the sites. Molecular concentrations and binding affinities decide the reaction propensities of individual binding. Once the model (molecules and rules) is defined, NFsim follows a kinetic monte carlo scheme [23], similar to the stochastic simulation algorithm (SSA) [24], to propagate the reactions. In this framework, the next reaction time and the next reaction event is determined in a stochastic manner. Importantly, NFsim does not require the predefinition of all molecular species (which would be impossible for polymerization), but rather generates new species on the fly. NFsim can make the distinction between the inter-complex binding and intra-complex binding which is very important for the current study and discussed in the result section (Fig. 3).

The simulations have units of molecular counts, but these can be readily converted to equivalent concentrations, which is how we present our results. We define the model using the BioNetGen Language (BNGL:http://bionetgen.org/) [25] and run multiple simulations using the UConn Health High Performance Computing Center. A typical 50 ms simulation of a system consisting of 9000 sites (1800 molecules) takes about 1 minute on a Xeon processor. The NFsim solver uses a random time step algorithm, based on the rate parameters, to optimize the simulation efficiency.

### Data Analysis

Once we execute multiple trials for a given simulation condition, for both SpringSaLaD and NFsim, we use custom Python scripts to collect and analyze the results across multiple trials. A “cluster” is a molecular network where each node is a multivalent molecule, and the physical bonds comprise the edges. We characterize the molecular networks to extract several biophysical properties of the system. A molecule with no bonds (edges) is considered to be “free” or a monomer. The outputs of SpringSaLaD and NFsim construct histograms of fraction of total molecules vs. number of molecules in a cluster (i.e. cluster size, where a monomer is a cluster of size 1) at each time point. We define a steady state after the time at which the distributions in the histograms fluctuate around a constant mean distribution, using a minimum of 50 trajectories.

Our statistical methods to analyze the cluster distributions would likely be useful to the biophysics community studying multivalent clustering and phase transition. We have organized and released the python code that is available at https://github.com/achattaraj/maximal_Solubility_product.

All the python packages are installed and managed by Anaconda distribution https://www.anaconda.com/products/distribution. Most frequently used packages are: Numpy (version 1.22.4), matplotlib (version 3.5.1) and pandas (version 1.4.3).

## Results

### The phase transition threshold is marked by the maximum solubility product

We first consider a pair of pentavalent molecules with different spacing between the binding sites (Fig. 1A). We quantify the clustering propensity of the system as a function of increasing molecular concentrations. Snapshots of the steady state distributions at 4 different concentrations are provided at the top of Fig. 1B. We checked that these systems were at steady state, not a kinetic metastable state, by running selected simulations for periods that were extended 10X the standard period (Fig. S3). Cumulative histograms of the cluster size distributions from 400 independent steady state time points are shown at the bottom of Fig. 1B. Each histogram displays the cluster size vs. fraction of total molecules within those clusters. The heavy-tailed nature of the distributions becomes more apparent at higher concentrations, with obvious bimodality at the highest concentrations. The average cluster occupancy or ACO is the mean of the distribution within each of these histograms. The ACO, in units of molecular counts, serves as a single parameter that measures the tendency of the molecules to form large clusters [14, 22]. The spatial proximity effect makes the “dimer” a special case, as seen by the prominent peak at cluster size 2 in all the histograms of Fig. 1B. We refer to this phenomenon as “dimer trap” which arises from the stoichiometry-matching effect, as described in our previous studies [22]. Wingreen and colleagues have used the term “magic number effect” [18, 26] to denote the stability of these stoichiometric complexes.

**Figure 1:**
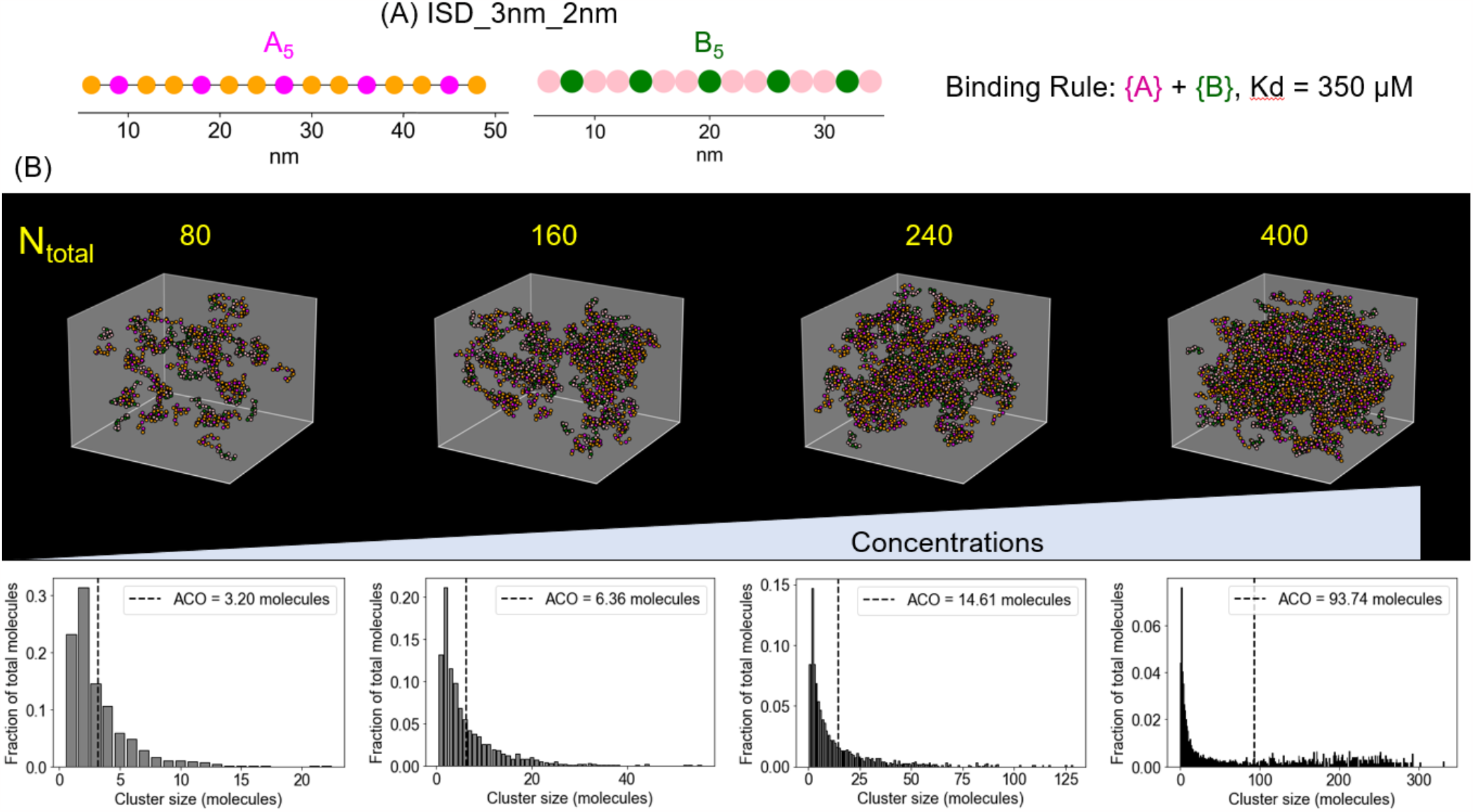
Quantification of multivalent molecular clustering. (A) SpringSaLaD representation of a pair of pentavalent binding partners. A_5_ contains 5 magenta binding sites and 10 orange linker sites (linear molecular length = 42 nm). Similarly, B_5_ contains 5 green binding sites and 10 pink linker sites (linear molecular length = 28 nm). For all sites, radius = 1 nm, diffusion constant = 2 µm^2^/s. For individual binding, dissociation constant, Kd = 350 µM (Kon = 10 µM^-1^.s^-1^, Koff = 3500 s^-1^). Simulation time constants, dt (step size) = 10^−8^ s, dt_spring (spring relaxation constant) = 10^−9^ s. (B) We randomly place N molecules of each type in a 3D reaction volume of 100*100*100 nm^3^. Then we titrate up N to increase the molecular concentrations (single steady state snapshots are displayed in the upper panel) and quantify the cluster size distributions at steady state (lower panel). The mean of the cluster distribution (dashed black line) is termed as average cluster occupancy or ACO. We collect molecular clusters over 4 post-steady state timepoints (10ms, 20ms, 30ms, 40ms) from each run across 100 trials. Therefore, ACO is computed over 400 independent realizations.

The cluster distributions in Fig. 1B appear to become more bifurcated with increasing concentration. However, this is best appreciated by examining individual steady state distributions rather than the cumulative histograms of Fig. 1B. Figs. 2A and 2B illustrate cluster distributions from single steady state time points in several individual runs at two different concentrations; importantly, the overall system fluctuates around a steady state at all of these times. At a lower concentration (Fig. 2A), molecules remain within a collection of small clusters. Although there are 120 molecules (60 of each type) in the system, we never see any cluster that is greater than 25 molecules within the sample of 42 individual histograms shown in Fig. 2A. To confirm that this behavior is not sensitive to the number of molecules in the system, we test the behavior with a larger system (Fig. S1); we place 200 molecules of each type (3.3 times the number in Fig. 2A) in a volume of 150*150*150 nm^3^ such that the molecular concentration is still ∼100 µM. We almost never see any cluster greater than 25 molecules although the total available in the system is now 400. However, as we increase the concentration individual large clusters do appear in the system. A higher concentration (Fig. 2B) shows strongly bifurcated cluster distributions where a single very large cluster coexists with multiple small oligomers. These individual large clusters are obscured in the cumulative histograms of Fig. 1B. Fig. S2 shows the steady state (last time point) cluster distribution of the system at these two concentration points for all 100 runs. This switch in clustering behavior from only small clusters to a distribution of small clusters coexisting with one or two large clusters is an indicator of the phase transition.

**Figure 2:**
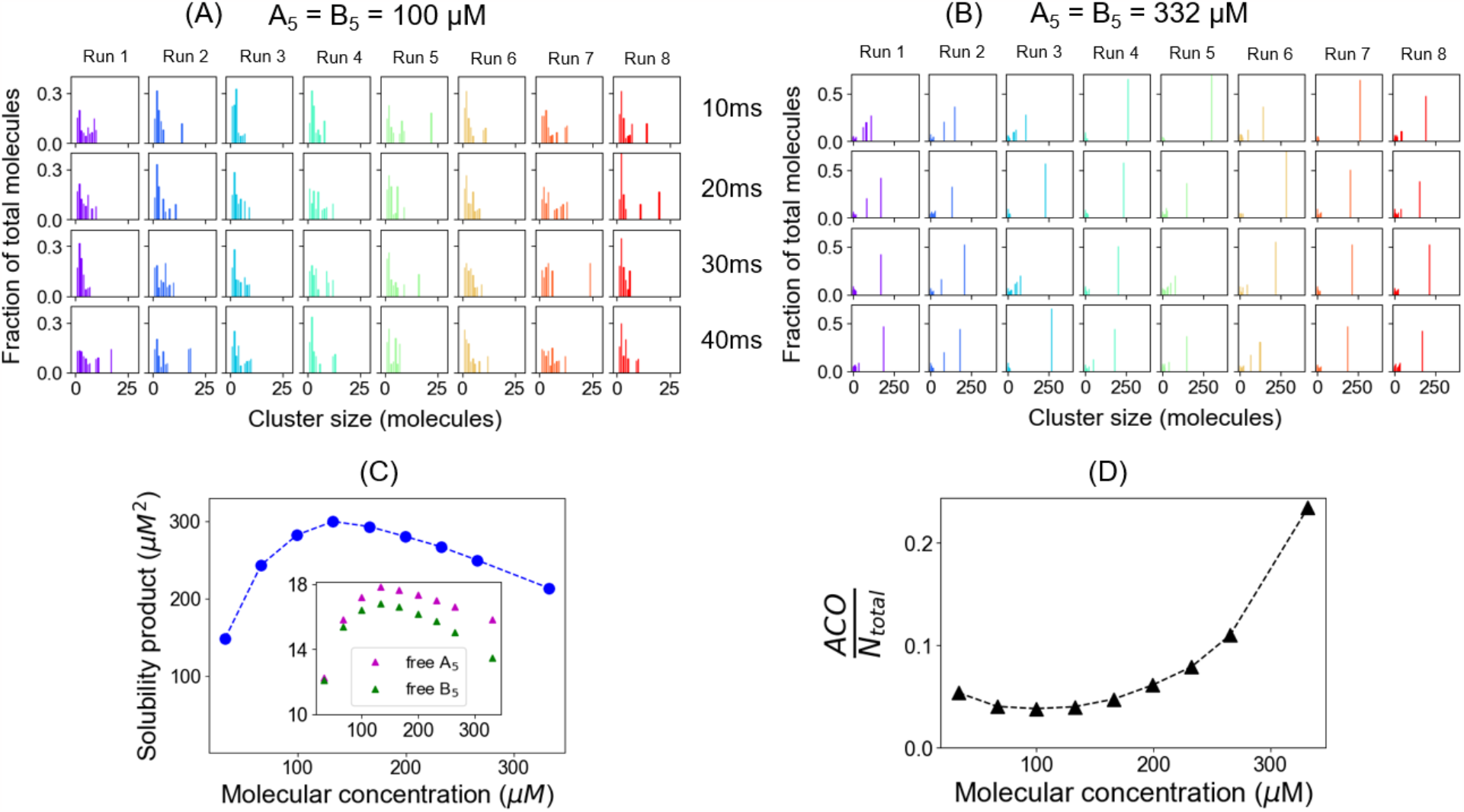
Solubility product maximum marks the phase transition threshold for a pair of pentavalent binders. (A, B) Time evolution of cluster distributions at a low (100 µM) and high (332 µM) concentration. For each concentration, 4 timepoints from 8 typical stochastic runs are displayed where each run is represented by a separate color. (C) Solubility product (SP) vs. molecular concentration of the pentavalent pair (illustrated in Figure 1A). We titrate up molecular counts in a fixed volume (100*100*100 nm^3^) and determine the monomer concentrations (free A_5_ and free B_5,_ inset) at steady state. For this system, N = [20, 40, 60, 80, 100, 120, 140, 160, 200] where N is the molecular count of each type. So, molecular concentration = [A_5_] = [B_5_]. We run 100 trials for each condition and sample 30 independent steady state timepoints for each trial. So, each free concentration point is an average over 3000 independent realizations. (D) Normalized average cluster occupancy (ACO) as a function of molecular concentration. ACO of the system is normalized by the total molecules (N_total_ = #A_5_ + #B_5_) present in that system. To quantify cluster distributions, we sample 4 timepoints (10ms, 20ms, 30ms, 40ms) from each run across 100 trials; so ACO is computed over 400 independent realizations.

**Figure 3:**
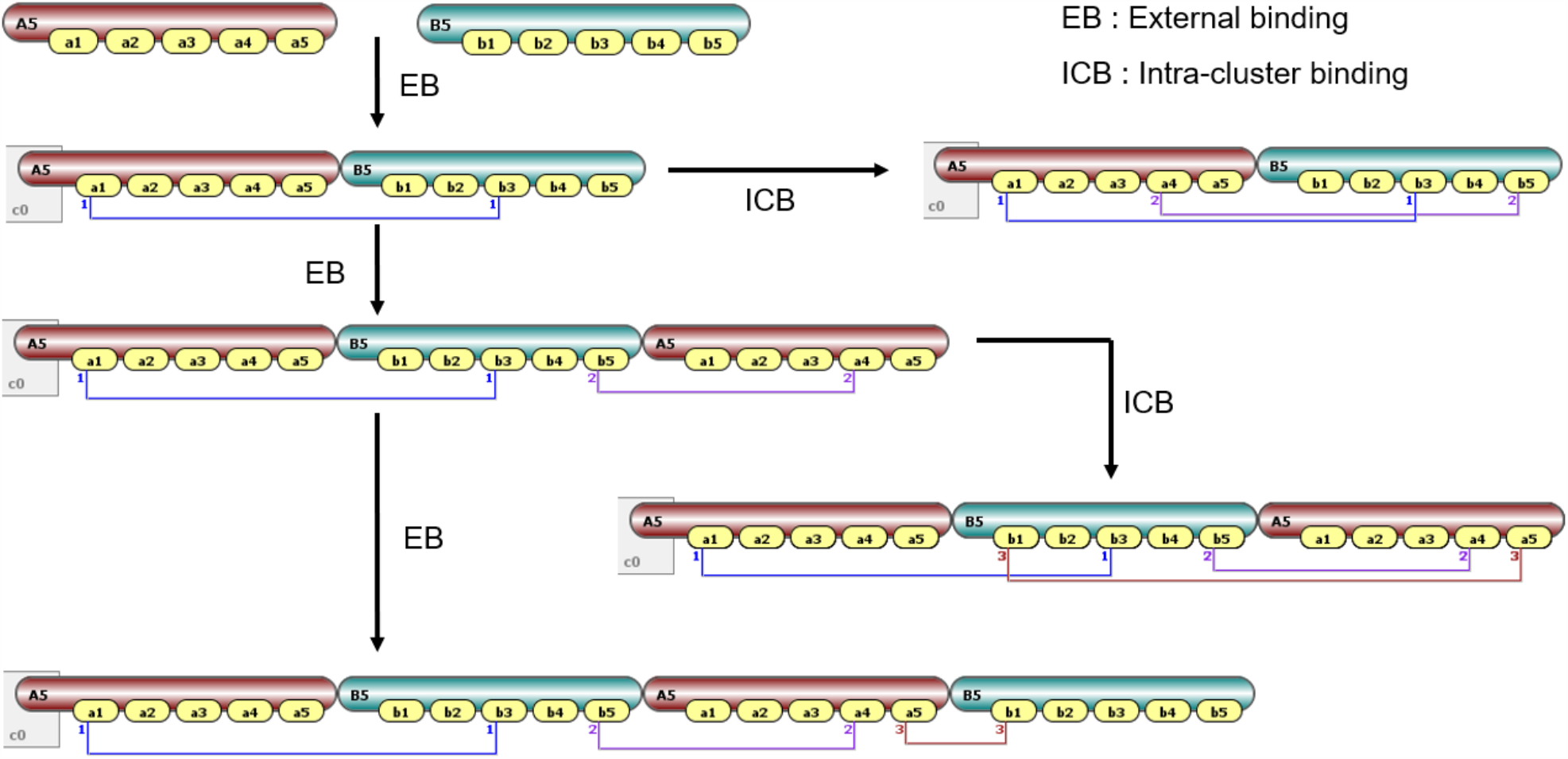
Illustration of binding rules for non-spatial NFsim model (Virtual Cell interface). When a molecular cluster grows by recruiting new molecules, it is called external binding (EB). When a new bond is established between two molecules that are already part of the same cluster, it is called intra-cluster binding (ICB).

The dynamics of clustering is also of interest; for example, in Run 3 of Fig. 2B, we see that the lone large cluster collapses completely at 30ms and another re-emerges at 40ms. Runs 5 or 6 demonstrate similar behavior. Fig. S3 indicates that the dynamic nature of clustering is preserved over longer timescales, even though the cluster distribution is heavily bifurcated. The dynamic cluster dissolution and formation could be important for the material properties of the cluster phase.

To explore the concentration threshold of the phase transition, we compute the solubility products (SP) (Fig. 2C) and ACO/N (Fig. 2D) as a function of total molecular concentrations. SP is simply the product of free monomer concentrations of A_5_ and B_5_. SP steeply increases up to a maximum and then drops gradually. Fig. 2C inset shows that concentrations of free A_5_ are always higher than free B_5_ at steady state, indicating that clusters recruit the shorter molecules B_5_ more than longer A_5_. Fig. S4 shows a SpringSaLaD simulation illustrating the conformational flexibility of the monomers; analysis of these simulations show that both end to end distance and radius of gyration is roughly 1.3 times higher for A_5_ than B_5_, which could account for the preferred recruitment of the smaller B_5_ molecules within the clusters. Importantly, at the same concentration as the SP maximum, ACO/N (Fig. 2D) displays a minimum. Before this point, the cluster size distribution has a shape that approximately decays exponentially (first 2 histograms in Fig. 1B) and ACO scales with the number of molecules entering the fixed volume. Beyond the minimum in Fig. 2D, the cluster size begins to build up faster than the total number of molecules (third and fourth histogram in Fig. 1B); in other words, as more molecules are fed into the system they are funneled into large clusters. As noted in our discussion of Figs. 2A and 2B, this threshold coincides with the appearance of lone large clusters in the individual trajectories. Thus, the minimum in Fig. 2D defines the concentration threshold for the phase transition. Comparing Figs. 2C and 2D shows that the maximum SP coincides with the phase transition threshold (∼ 133 µM) of the system.

We also checked whether the finite size of the system has any siginificant effect on the SP profile. Figure S5 compares two scenarios where the system size differs by a factor 4, i.e., smaller systems have 1x molecules in the nominal volume, while the larger systems have 4x molecules in 4 times the nominal volume. In other words, we maintain concentrations for two different sized systems and compute the SP profiles. Gratifyingly, both profiles point towards the same total concentration where SP reaches a maximum, that is, the phase transition threshold is the same for both systems and is therefore independent of the system size. We also notice that the SP curve is slightly upshifted for the larger system. This can be attributed to a boundary effect at the edges of simulation box: the smaller system has a larger surface-to-volume ratio which increases the effective concentrations by an excluded volume effect with the hard spheres at the boundaries. The higher effective concentration slightly shifts the curve down for the 1x case. Importantly, however, Fig. S5 allows us to conclude that the maximum SP displays its maximum at the same total concentration irrespective of the number of molecules (i.e. volume) of the system.

### Intra-cluster binding shapes the solubility product profiles

We previously reported [14] that the onset of multivalent biomolecular condensate formation is approximately characterized by a solubility product constant (Ksp) where a relatively flat SP is maintained beyond the phase transition threshold, funneling the additional molecules to the cluster phase. In that previous study, we used the non-spatial NFsim [15] solver to deduce the Ksp principle, where we could isolate the effects of valency from the structural features of the molecules (e.g., NFsim does not account for steric effects, molecular flexibility and intra-cluster site physical proximity). Since NFsim does not track the spatial proximity of the binding sites, our prior study permitted only external binding (EB, Fig. 3); this was done to prevent purely statistical crosslinking that could arise from intra-cluster binding (ICB, Fig. 3) unconstrained by steric effects or physical proximity. A spatial solver like SpringSaLaD directly accounts for proximity and steric effects, permitting us to explore the Ksp concept in more realistic models. Indeed, Fig. 2C shows that beyond the threshold concentration the SP starts to dip down instead of staying constant, indicating a positive feedback for the growth of clusters. However, before further exploring the influence of proximity and steric effects with SpringSaLaD, we sought to assess how only the statistical effect of intra-cluster binding impacts the distribution of clusters as a function of concentration; this could be achieved by comparing NFsim calculations with and without intra-cluster binding enabled.

We consider the clustering of a pentavalent binding pair in the presence or absence of intra-cluster binding (Fig. 4). Fig. 4A displays how intra-cluster binding affects the SP profile. Interestingly, the maximum SP is reached at the same total concentration with or without intra-cluster binding. In other words, the phase transition threshold is ∼30 µM for both systems. However, beyond the threshold, SP goes down gradually with intra-cluster binding much like the SpringSaLaD simulations in Fig. 2C. The clustering behavior also differs (Figs. 4B, 4C and 4D). ACO increases more dramatically with intra-cluster binding (Fig. 4B), corresponding to a sharper phase transition. The absence of intra-cluster binding puts an upper limit to the bound fraction of clusters. Fig. 4C shows that maximal bound fraction is 0.4 for the pentavalent pair both below (30µM) and well above (90µM) the phase transition, which means that only 2 out of 5 sites can be occupied on average without intra-cluster binding. When we analyze the steady state bond count distribution per single molecule (Figs. 4E, 4F), we notice that the mean of the distribution always remains less than 2 bonds per molecule (blue bars) without intra-cluster binding. With intra-cluster binding, beyond the threshold (90 µM), cluster distribution is more bifurcated (Fig. 4D, lower panel) and a higher number of sites are now occupied (bound fraction = 0.46). The patterns of bound fraction profiles are also different in the presence of intra-cluster binding, as additional crosslinkings (ring structures) are now permitted. With intra-cluster binding, the bond count distribution yields an average of 2.24 bonds per molecule (bound fraction = 2.24/5 ∼ 0.45) above the threshold.

**Figure 4:**
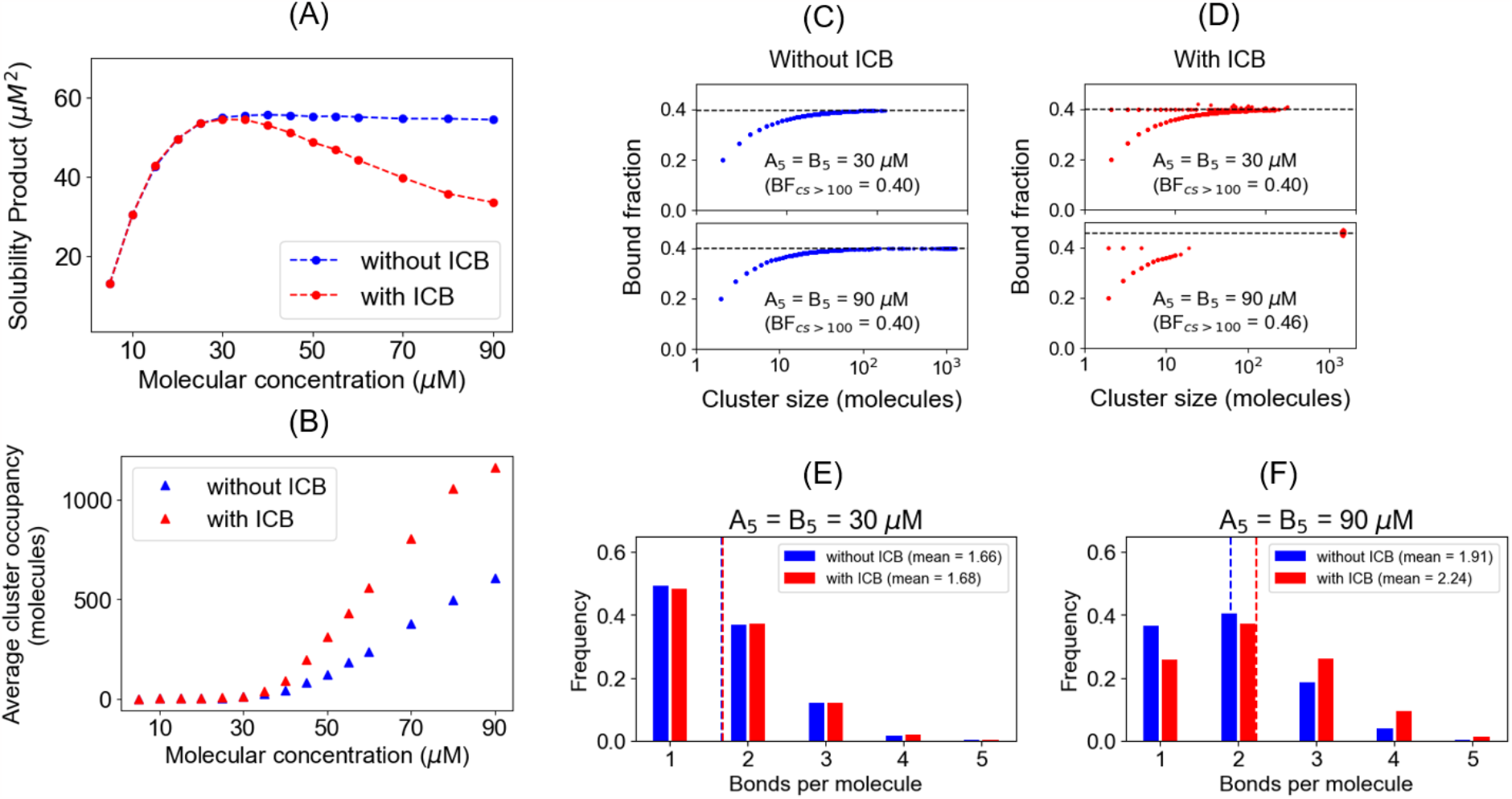
Intra-cluster binding (ICB) causes a gradual drop in solubility product by promoting higher degree of crosslinking. (A) Solubility product profiles with and without the ICB. Each concentration point is an average over 2000 steady state points (20 timepoints X 100 runs). We titrate up the molecules in a fixed volume to increase the concentration. Molecular counts of each type = [50, 100, 150, 200, 250, 300, 350, 400, 450, 500, 550, 600, 700, 800, 900]. Dissociation constant = 350 µM. (B) Quantification of clustering with and without the ICB. Cluster distributions are averaged for the last timepoint of 100 runs. (C, D) Binding saturation of clusters near (30 µM) and above (90 µM) the phase transition threshold, without and with the ICB respectively. The bound fraction is the ratio of bound sites to total sites within a cluster. The dashed line represents the limiting value that the large clusters (> 100 molecules) converge to. Note the logarithmic scale of the abscissa. (E, F) Quantification of molecular crosslinking with and without the ICB, near (30 µM) and above (90 µM) the phase transition threshold respectively. The dashed line denotes the mean of the bond count distribution.

The clustering patterns of non-spatial (NFsim) and spatial (SpringSaLaD) systems are quite similar when appropriately scaled. To compare the SP vs. concentration curves, in Fig. 5 we normalize the axes of solubility product and concentrations (equivalent to comparing non-dimensionalized variables). When SP reaches a maximal level (SP_max_), the associated concentration is set as the threshold (C_threshold_). We then normalize the SP axis with SP_max_ and concentration axis with C_threshold_ (Fig. 5A). Remarkably, we find that SpringSaLaD SP profile (green triangles) coincides with the NFsim (with intra-cluster binding) SP profile. When we analyze the clustering trends for these systems (Fig. 5B), NFsim (with intra-cluster binding) slope is much higher than SpringSaLaD, beyond the threshold. However, NFsim (without intra-cluster binding) system shows a less pronounced difference with SpringSaLaD. Since spatial effects are approximated in SpringSaLaD and ignored in NFsim, we may consider the SpringSaLaD model output to be a better biophysical picture of the system. However, Fig. 5 suggests that NFsim with intra-cluster binding does capture the partitioning of monomers. For the cluster distributions, NFsim without intra-cluster binding seems to yield a better trend, most likely by avoiding the non-physical cross-links. The entropic cost of clustering is always higher in spatial system as the non-spatial system cannot account for spatial degrees of freedom (translation, rotation/conformation etc.). This gets reflected in the lower normalized ACO for SpringSaLaD compared to both NFsim systems (Fig. 5B). As noted above, the spatial proximity effect favors stoichiometry matching [22], producing a “dimer trap” (Fig. 1B), which is absent in the non-spatial NFsim simulations.

**Figure 5:**
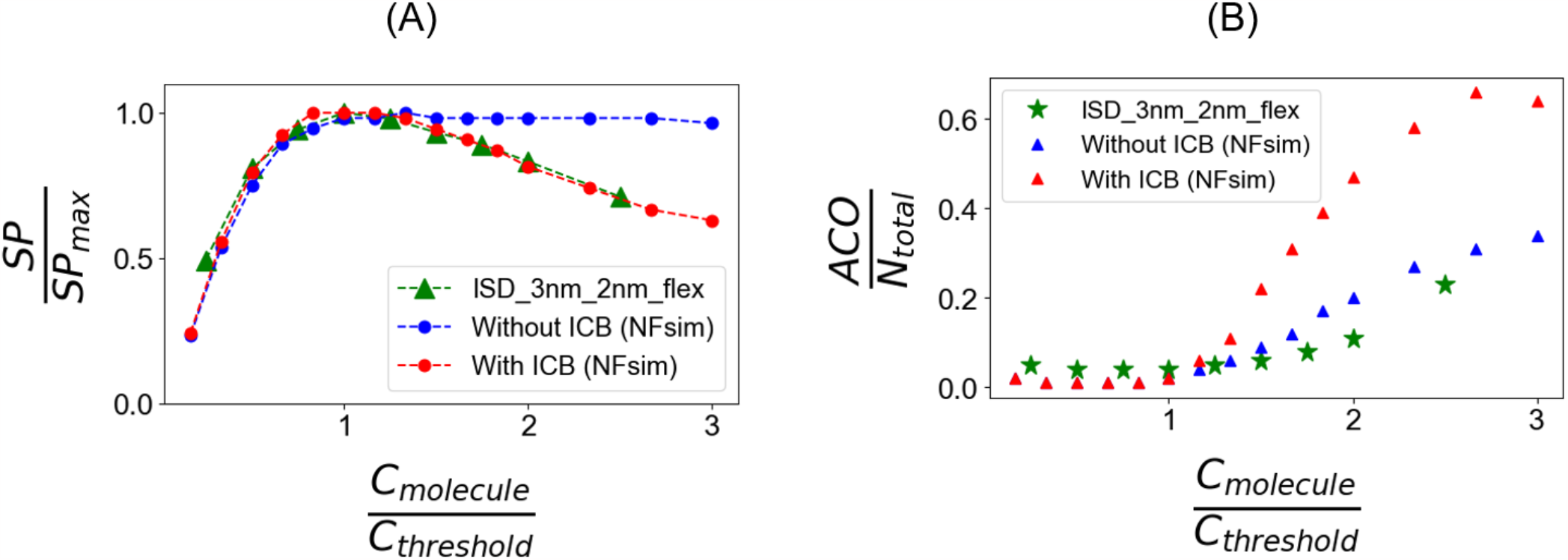
Comparison of molecular partitioning between free and cluster state using spatial and non-spatial simuators. (A) Normalized solubility product profiles. When the solubility product (SP) reaches the maximum (SP_max_), the corresponding concentration is called threshold (C_threshold_). The horizontal axis is normalized by threshold concentration and the vertical axis by maximal solubility product. (B) Normalized ACO as a function of normalized concentrations. The SpringSaLaD structure from Figure 1A is used here to compare with NFsim results.

To check the generality of the maximal solubility product principle, we next study the mixed valent systems in Fig. 6. We first consider a two component system: A_5_– B_3_ (Fig. 6A) where the relative total concentrations of the molecules are kept at 3 : 5 to adjust for the binding stoichiometry. For this mixed-valent system, SP = [free A_5_]^3^ * [free B_3_]^5^ [14]. Consistent with the simpler system in Fig. 4, the solubility product reaches the same maximal point in the absence (blue) and presence (red) of intra-cluster binding (upper panel, Fig. 6C). Beyond that point, SP starts to drop gradually, accompanied by a sharper increase in clustering (lower panel, Fig. 6C). We plot the SP and ACO as a function of total concentration (A_5_ + B_3_, which are different for this mixed-valence system); the threshold for phase transition still corresponds to the maximal SP. When we consider a three-component mixed-valence system (Fig. 6B), the behavior (Fig. 6D) is remarkably similar to the 2 -component systems, whether matched valence (Fig. 4) or mixed valence (Fig. 6C). For 6D, SP = (free A_3_)^2^ * (free B_1_3_)^6^ * (free C_6_)^3^. In all these cases, a maximal solubility product marks the threshold beyond which the ACO starts to go up sharply, indicating a phase transition.

**Figure 6:**
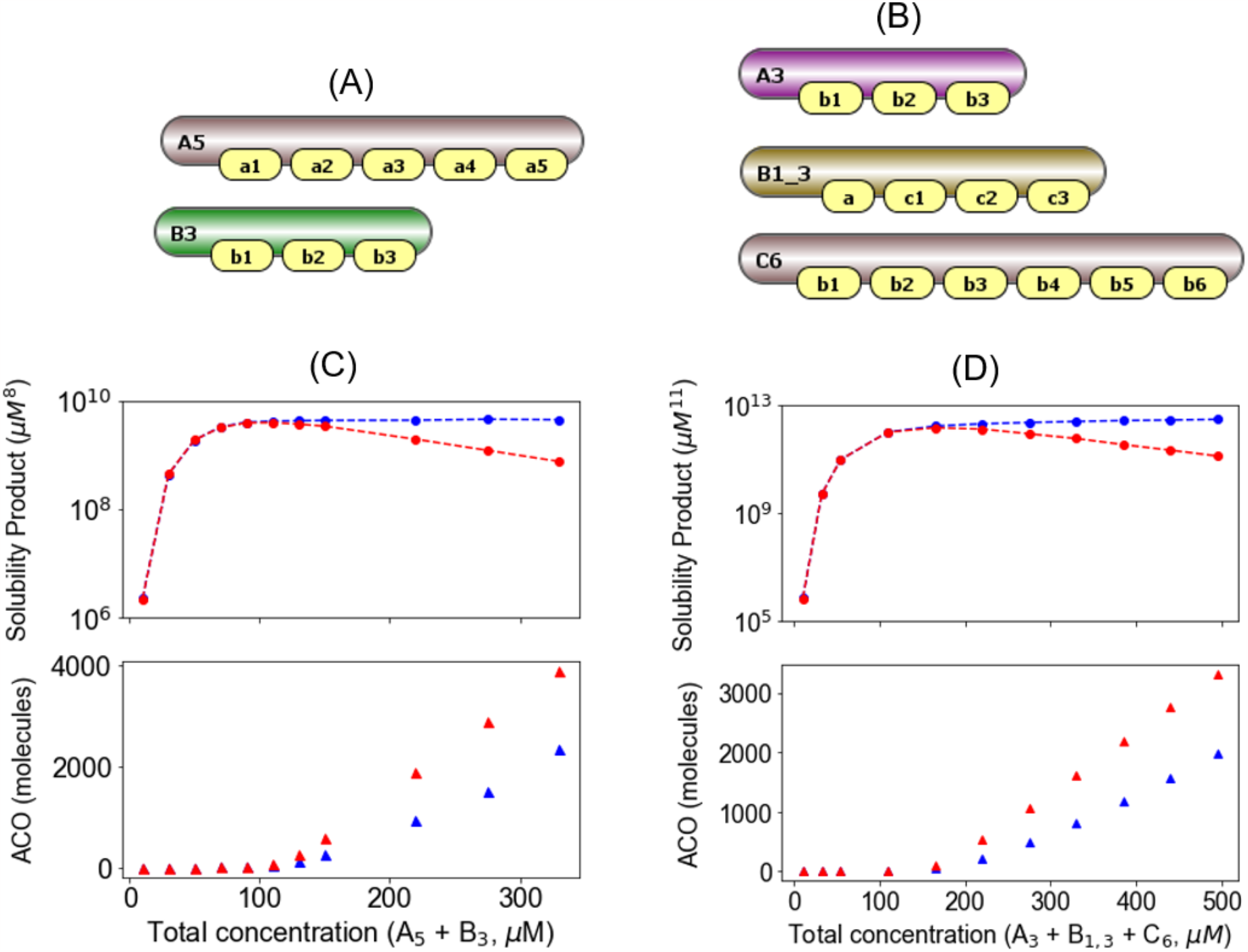
Maximal solubility product yields the phase transition threshold for mixed-valence systems. (A) A pair of pentavalent and trivalent molecules. Each A site can bind to any of the B sites with an affinity of 350 µM. (B) A three-component system where B_1_3_ can bind to both A_3_ and C_6_. Name of the sites reflect their complementary binding sites; for example, B_1_3_ has one binding site for A_3_ and three binding sites for C_6_. All pairwise interactions are assumed to have an affinity of 350 µM. (C) Solubility product (upper panel) and ACO (lower panel) profiles with (red) and without (blue) the ICB for A_5_ – B_3_ pair. Each concentration point is an average over 2000 steady state points (20 timepoints X 100 runs). We titrate up the molecules in a fixed volume to increase the concentration. To account for the binding stoichiometry, A_5_ : B_3_ is maintained at 3:5 ratio. A_5_ counts = [ 60, 180, 300, 420, 540, 660, 780, 900, 1320, 1650, 1980] and B_3_ counts = [ 100, 300, 500, 700, 900, 1100, 1300, 1500, 2200, 2750, 3300]. For this mixed-valent system, SP = (free A_5_)^3^ * (free B_3_)^5^. (D) Solubility product (upper panel) and ACO (lower panel) profiles with (red) and without (blue) the ICB for the three components system. Molecular counts are titrated up maintaining a ratio of A_3_ : B_1_3_ : C_6_ = 2 : 6 : 3. A_3_ counts = [ 20, 60, 100, 200, 300, 400, 500, 600, 700, 800, 900]; B_1_3_ counts = [ 60, 180, 300, 600, 900, 1200, 1500, 1800, 2100, 2400, 2700] and C_6_ counts = [30, 90, 150, 300, 450, 600, 750, 900, 1050, 1200, 1350]. For this three-component system, SP = (free A_3_)^2^ * (free B_1_3_)^6^ * (free C_6_)^3^.

### Interplay of molecular length, flexibility and steric effect controls phase transition

We next set out to dissect the role of molecular structural motifs on the phase transition propensity. We use multiple structural pairs (Fig. 7) to study the effects of molecular lengths, flexibility and steric effects. Structures in Figs. 7A and 7B (along with Fig. 1A) enable us to probe the effects of molecular length and relative inter-site distances between binding partners. We also create a rigid version of the molecules by placing an end-to-end link, which serves as a distance constraint between the terminal sites. Fig. 7C depicts a binding pair that contains bulkier linker sites to probe for steric effects.

**Figure 7:**
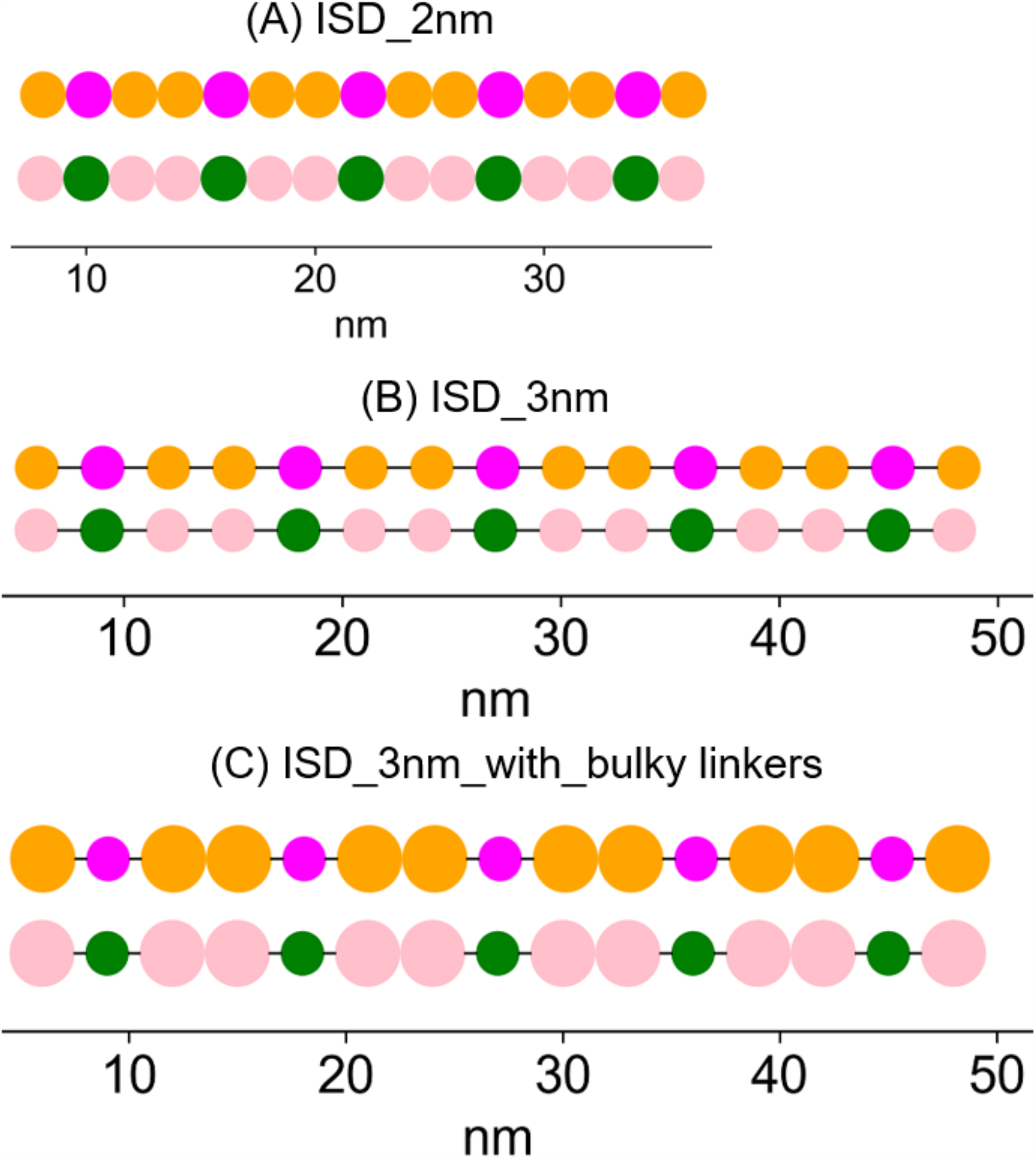
SpringSaLaD models to probe the effect of molecular structures on solubility product profile. (A) Pair of pentavalent binders with inter site distance (ISD) = 2 nm. Linear molecular length, A_5_ = B_5_ = 28 nm. (B) Pair of pentavalent binders with inter site distance (ISD) = 3 nm. Linear molecular length, A_5_ = B_5_ = 42 nm. For all sites in structures A and B, radius = 1 nm. (C) Linear molecular lengths are same as B, except for bulkier linker sites. Radius, binding (magenta and green) sites = 1 nm and linker (orange and pink) sites = 1.5 nm. Diffusion constant and binding rate constants for structures A, B and C are same as the pentavalent binders specified in Figure 1A.

Figs. 8A summarizes the SP profiles of five structural pairs having different molecular lengths and flexibility, including also the original “ISD_3nm_2nm_flex” discussed in relation to our earlier results. Importantly, comparing Fig. 8A and 8B extends the generality of Figs. 2, 4 and 6, showing how in all cases the concentration at which the maximum SP is reached is also the concentration at which the ACO/N begins to grow. Thus, the maximum SP appears to universally mark the concentration threshold for phase transition. Fig. 8C and 8D display the normalized SP and normalized ACO respectively. Most of the structures show remarkable similarity in these normalized plots.

**Figure 8:**
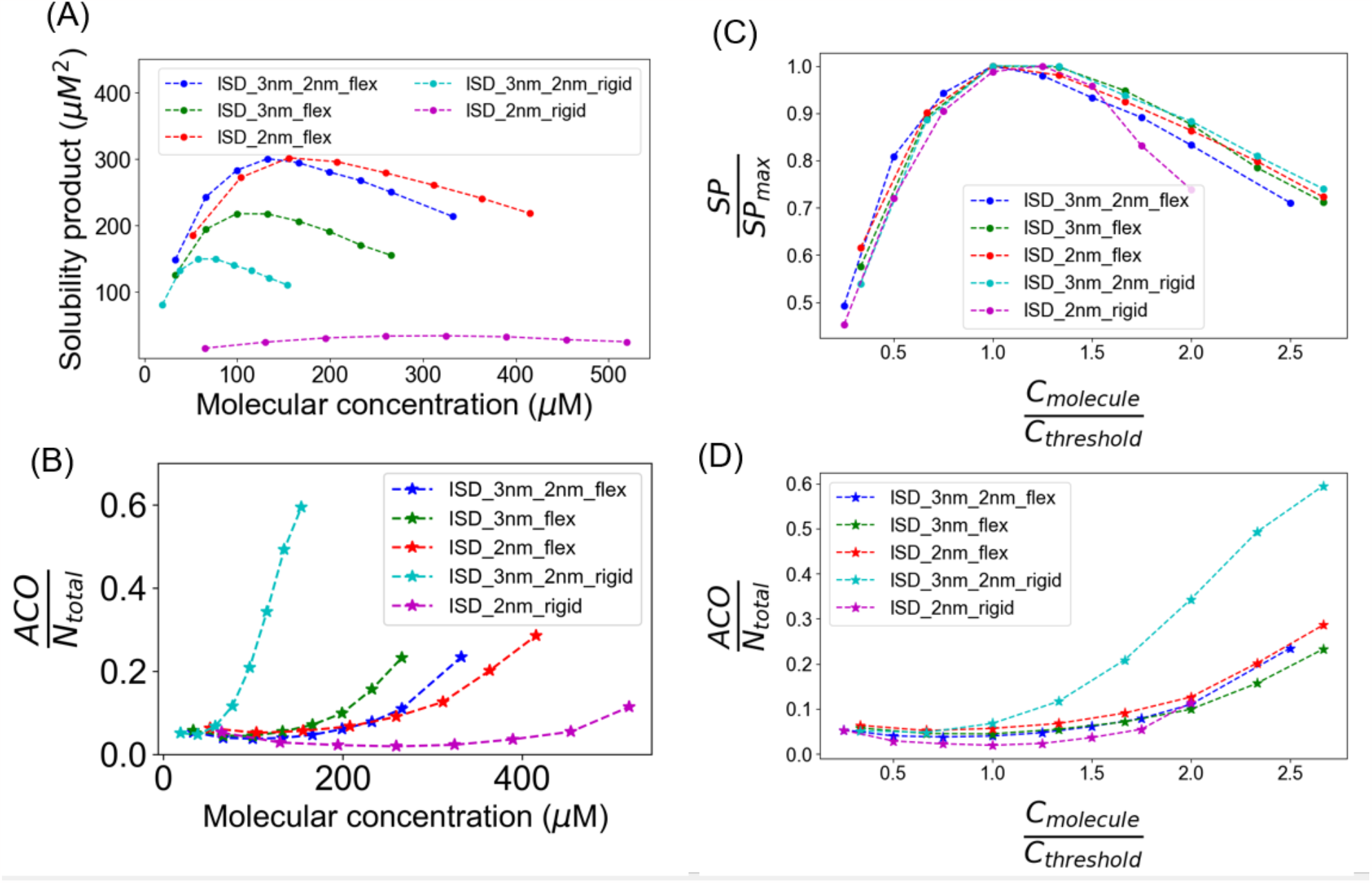
Interplay of distance mismatching and flexibility controls phase transition. (A) Solubility product profiles for five structural systems. The legends indicate the structures of the molecular pairs, referring to Figs.1A, 7A and 7B. The respective rigid pairs are obtained by creating an additional link between the terminal sites of each molecule, which works as a constraint of end to end distance. Reaction volume, ISD_3nm_2nm_flex = ISD_3nm_flex = 100*100*100 nm^3^ ; ISD_2nm_flex = 80*80*100 nm^3^; ISD_3nm_2nm_rigid = 120*120*120 nm^3^; ISD_2nm_rigid = 80*80*80 nm^3^. Except for ISD_3nm_2nm_flex (described in Figure 1B legend), molecular count of each type = [20, 40, 60, 80, 100, 120, 140, 160]. We run 100 trials for each condition and sample 30 steady state timepoints for each trial. So, each free concentration point is an average over 3000 steady state points. (B) Normalized ACO as a function of molecular concentrations. Clusters are sampled over 400 steady state points. (C, D) Normalized solubility product and Normalized ACO profiles as a function of normalized concentrations.

Noting especially that a higher SP indicates a lower overall thermodynamic tendency to form clusters, a more detailed examination of Fig. 8, suggests the following:

1. Flexible molecules (blue, red, green) have higher SP compared to the rigidly extended ones (cyan and magenta). This finding, which has also been pointed out previously [22, 27], is consistent with the idea that more extended structures will keep binding sites exposed and available.
2. Longer flexible pair (green, ISD_3nm_flex) has lower SP than flexible pairs with shorter (red, ISD_2nm_flex) or mixed lengths (blue, ISD_3nm_2nm_flex).
3. Except for the ISD_2nm_rigid case, all SP profiles appear very similar in normalized SP vs normalized concentration plane (Fig. 8C).
4. Except for ISD_3nm_2nm_rigid system, normalized ACOs are very similar to each other.

As best appreciated by examining Fig. S6, these results reflect an interplay between distance matching between binding sites and the flexibility of linker regions. For flexible molecules, the dimer trap is always visible (Figs. S6A, S6C) irrespective of distance matching between binding sites in partner molecules. However, for rigid structure, distance-matched partners display very prominent dimer traps (Fig. S6D) while distance-mismatched partners are unable to form multiply-bound dimers and form really large clusters (Fig. S6B). Since rigid structures have lower initial entropy, cross-linking is more favorable (cyan plot in Figs. 8B and 8D). Interestingly, this clustering behavior happens to be very similar to the NFsim (with intra-cluster binding) simulations (Fig. 5B).

For distance-matched rigid structures (ISD_2nm_rigid), the dynamics for approach to steady state (Fig. S7D and Video 1) is also distinct from systems that can largely avoid the dimer trap: it starts with a highly cross-linked state (large ACO) and gradually relaxes to a lower steady ACO value with a dominant dimer population. The clustering dynamics for the other systems behave very differently (Figs. S7A, S7B, S7C) where the dimer trap is not severe. This analysis supports the role of both rigidity (as noted in [26]), and binding site distance matching in the operation of the dimer trap. Such structural features will play a key role in distinguishing between the assembly of strictly stoichiometric complexes which we have called molecular machines [28]) vs. non-stoichiometric condensates (which we have called pleomorphic ensembles [28, 29]).

Next, we explore the effects of bulky linker sites (“ISD_3nm_with_bulky linkers” in Fig. 7). Figs. 9A and 9B show that binders with bulkier linkers (higher steric effects) have very high SP and lower ACO compared to the system with smaller linkers (lower steric effect), indicating lower tendency for phase transition. Gratifyingly, once again the maximum SP coincides with the minimum ACO/N_total_. Interestingly, the drop in normalized SP (Fig. 9C) is steeper for the bulky linkers (although normalized ACO runs below (Fig. 9D) the lower steric system). Clusters composed of bulkier molecules would more effectively trap monomers causing a steeper drop in SP.

**Figure 9:**
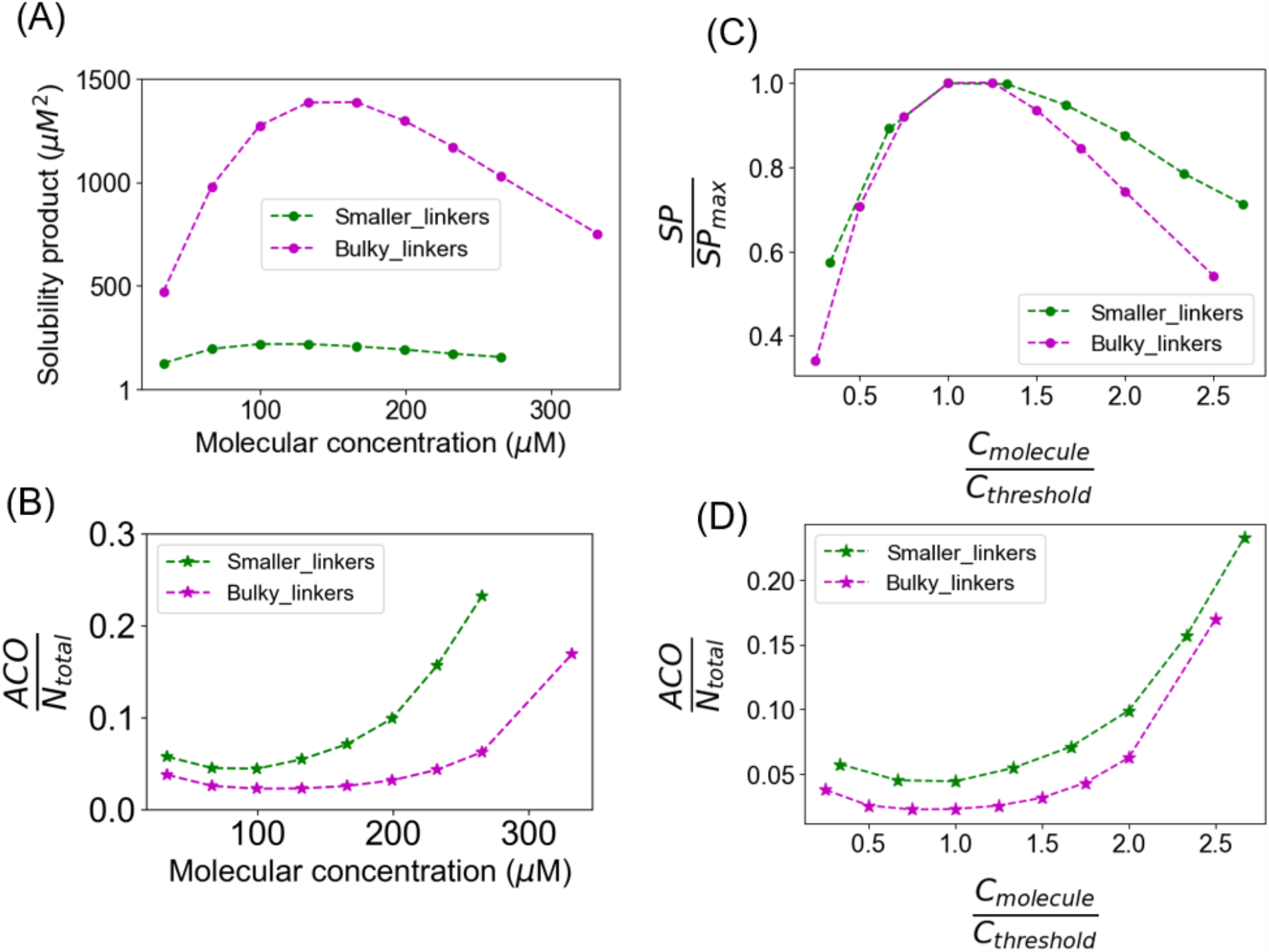
Steric hindrance reduces phase transition propensity. (A) Solubility product profiles for systems 7B (ISD_3nm_smaller_linkers) and 7C (ISD_3nm_bulky_linkers). For bulky_linker system, reaction volume = 100*100*100 nm^3^, and molecular count of each type = [20, 40, 60, 80, 100, 120, 140, 160, 200]. We run 100 trials for each condition and sample 30 steady state timepoints for each trial. So, each free concentration point is an average over 3000 steady state points. (B) Normalized ACO as a function of molecular concentrations. Clusters are sampled over 400 steady state points. (C, D) Normalized solubility product and Normalized ACO profiles as a function of normalized concentrations.

### Biophysical properties of clusters correlates with structures of component molecules

Beyond assessing how structural features of the binding partners influence the threshold for large cluster formation, we also sought to see how they influence the clusters themselves. We therefore analyze the SpingSaLaD output to extract some interesting biophysical properties. Fig. 10A displays the radius of gyration trend as a function of cluster size, for four flexible systems. For each cluster, we compute a centroid and then a radius of gyration around that. As might be expected, clusters composed of shorter molecules (red points) are, on average, more compact than longer or mixed molecules (green or blue points). Clusters composed of bulkier molecules (Fig. 7C) are the largest. The radius of gyration of the monomers (Fig. S8A) exactly follow the same trend. This raises a question about the connectivity of molecules inside the clusters. The intra-cluster connectivity should inversely be related with the degree of compaction. When we quantify the degree of cross-linking (Fig. 10B), we indeed find that bulky molecules are less cross-linked than other systems. Higher connectivity in shorter binders (ISD_2nm) results in more compact clusters, while the other two systems (ISD_3nm and ISD_3nm_2nm) fall in the middle.

**Figure 10:**
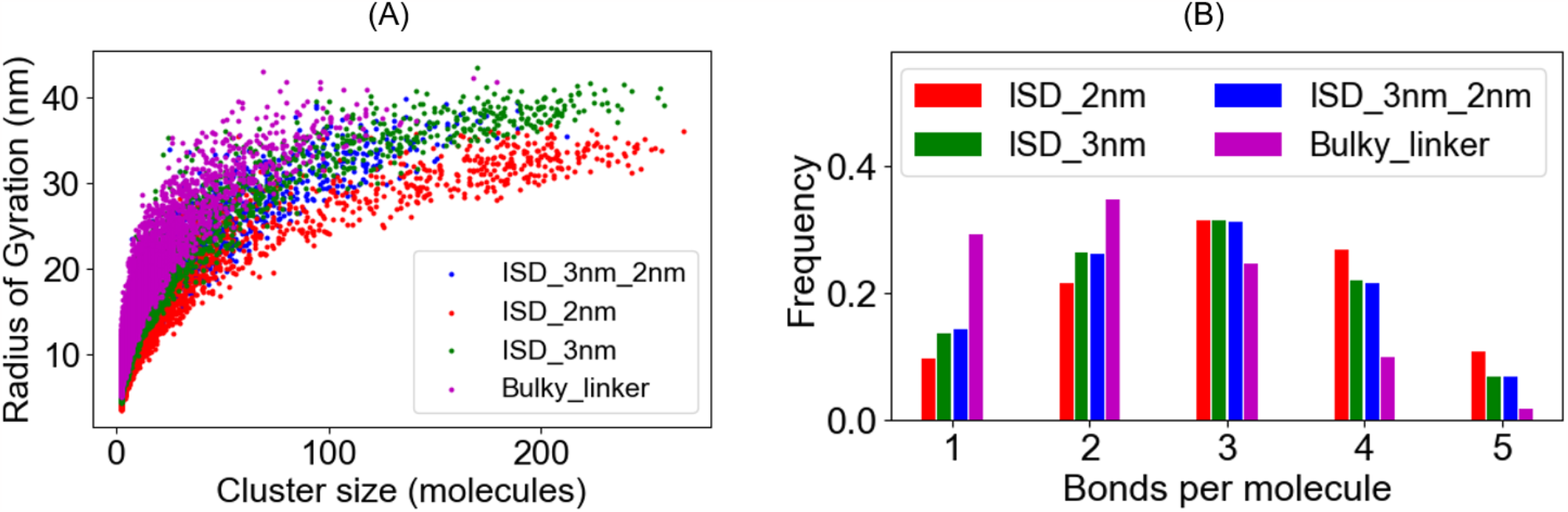
Biophysical properties of clusters depend on the structures of component molecules. (A) Cluster specific radius of gyration and (B) Bond count distribution for four structural pairs. For each system, total molecular counts = 160 + 160 = 320. Clusters are collected over 4 timepoints (10ms, 20ms, 30ms, 40ms) across 100 trials; total realizations = 400. All the component molecules are flexible.

Rigid structures (Fig. S8B) are expected to form less-compact clusters. Fig. S9A confirms that the ISD_3nm_2nm_rigid molecules create a larger network with less compaction. The expanded configurations of clusters composed of rigid structures offer more binding opportunity compared to the compact flexible counterpart, allowing many more monomers to be recruited into the network (cf., cyan curves in Figs. 8A and B). This also accounts for the difference in bond-count distributions, which show that clusters composed of rigid molecules have relatively fewer bonds per molecule (i.e., less intra-cluster bonds) as shown in Fig. S9B.

## Discussion

Clustering of multivalent biomolecules underlies the formation of biomolecular condensates, characterized by a transition from a homogeneous dispersed phase to a heterogenous clustered phase. The physics of single component (homotypic) condensation is relatively simple, where condensates appear beyond a threshold molecular concentration and leave the threshold dilute phase concentration unchanged [13]. In other words, the invariance of dilute phase concentration marks the phase transition boundary of such homotypic systems. However, due to binding stoichiometry and other biophysical constraints, multi-component (heterotypic) condensates display more complex behavior. Looking at titration curves of individual molecular components does not yield a homotypic-like threshold boundary [30]. This inapplicability of a simple concentration threshold requires more complex theories to describe the formation of heterotypic condensates. Multiple theoretical and computational efforts [31, 32] have attempted to address the issue with the concept of “tie-lines” where the slope of the tie-line is determined by the interplay of homotypic vs heterotypic interactions. We previously reported a simple approximation inspired by the ionic equilibrium of salt solutions. Using primarily non-spatial stochastic network free simulations, we proposed that formation of heterotypic condensates may be described by a solubility product (SP) [14] where the SP is defined as the stoichiometry-adjusted product of the dilute phase concentrations of the unbound constituent molecules. This SP converges to an approximate constant level when the system starts to form large clusters.

The results reported in this work help further refine the solubility product concept using spatial Langevin dynamics simulations to show that the maximum SP marks the concentration threshold of the phase transition, independent of the varying molecular structures and binding stoichiometries explored in this study (Figs. 2, 4, 6, 8 and 9). This is demonstrated by a coincidence of the maximum SP with the concentration at which the clusters grow more rapidly than the increase in available molecules (i.e. ACO/N_total_ begins to increase) for these various structural features (Figs. 2, 4, 6, 8 and 9). Our work here only explored stoichiometric concentrations of binders. As shown in our previous work [14], titration of non-stoichiometric concentrations of multivalent binders also reveals a maximum SP that marks the threshold for formation of large clusters. Importantly, the maximum SP in the non-stoichiometric cases are always below the maximum SP for the stoichiometric titrations.

The intra-cluster binding (intra-cluster binding) or ring structures strongly drive the clustering process beyond the threshold (Fig. 4). Therefore, beyond the threshold, SP gradually drops due to a gradual reduction in monomer dissociation from the highly cross-linked clusters. Using a different non-spatial solver, Falkenberg et al. [29] previously showed that the percolation threshold of multivalent binders can be robustly identified with the onset of ring formation (i.e. intra-cluster binding). Our current analysis is consistent with that finding. Beyond the threshold, more molecules cluster together and intra-cluster binding statistically becomes more operative to introduce more crosslinks. With higher degree of crosslinking, the probability of monomers coming off the clusters goes down; that is, the effective dissociation constant reduces. This post-threshold reduction of monomer release results in the drop of monomer concentrations, hence SP decreases.

Normalized SP (SP/SP_max_) and ACO (ACO/N_total_) are important and useful dimensionless parameters that enable us to compare systems with different biophysical attributes. Below the threshold for phase transition, the slope of the ACO/N_total_ vs. concentration curve is roughly 0 or slightly negative, meaning that the cluster size distribution does not grow with the number of molecules available in the system (as also shown in our previous work [14, 22]). Above the threshold, ACO/N_total_ has a positive slope and the slope goes up with higher concentrations; our spatial simulations are limited in size, but based on our previous work [14, 22], as we scale up the system size (i.e. increasing N), clusters will grow to the micron-scale at the macroscopic limit. What is remarkable here is how invariant the shapes of these plots of concentration vs. SP or ACO are when they are all non-dimensionalized (Figs. 5, 8, 9.). For SP/SP_max_, the curves are all similar below the threshold, and most are also aligned above it. These similarities suggest that the statistical probabilities of binding that solely drive clustering in the non-spatial systems (NFsim with intra-cluster binding), are sufficient to control the patterns of clustering in the spatial systems as well (Fig. 5A, 8C). However rigidified structures with bulky linkers (Fig. 9C) do show more negative slopes above the phase transition; once they are formed, these structures produce more sterically confining condensates, accounting for the (relatively) lower concentrations of free monomers in equilibrium with the clusters. ACO/N_total_ measure the tendency to form large clusters and when plotted against concentration normalized to the threshold concentration, the curves are again remarkably similar. The primary exception (Fig. 8D, cyan curve) is the case of rigid pentavalent binding pairs where the distances between binding sites are mismatched between the 2 molecules (the rigid version of Fig. 1A). We can understand the greater tendency to cluster for this system because the rigid structure lowers the entropy of the monomers so that less entropy is lost upon clustering; furthermore, that the binding site distances are not matched, prevents the “dimer trap” that limits the growth of clusters in the rigid matched “ISD_2nm_2nm” system (compare Figs. S6B and S6D.) Distance matched rigid molecules will always form stochiometric complexes (dimer for two-component homovalent system); this is likely a recipe to build “molecular machines” (for example, ribosome, proteasome etc.) where all the components fit together to create an interlocked structure. Flexible molecules (proteins with disordered region, for example), on the other hand, will form non-stoichiometric clusters (condensates) as the weak interactions are not strong enough to overcome the entropic effects.

It remains to be seen how well our work can predict biophysical properties of these systems. We have been inspired to devise our simulations based on a pair of seminal experimental papers [30, 33], which carefully analyzed, *in vivo* and *in vitro*, respectively, the ability of heterotypic systems to phase separate as a function of the total concentrations of their constituent molecules. We were able to show previously [14] that our approach qualitatively explains the patterns uncovered by these experiments. We may also be able to predict material properties of condensates such as viscosity or intra-cluster diffusion. Cluster radius (Fig. 10A) scales differently for different molecular structures. Interplay of flexibility and steric hindrance, imparted by the linker regions, controls the extent of cluster compaction. Weak non-specific interactions including homotypic interactions may play additional important roles in determining the compaction once molecules are assembled inside a cluster via specific interactions [34]. It would be important to consider these to see if some of the conclusions of this work might be further generalizable. Viscoelastic properties of condensates are a function of degree of molecular crosslinking [35, 36]. Towards that end, the bond count distribution (Fig. 10B) within clusters provides pertinent information. Finally, while we define SP in terms of monomer concentrations, the simulations both in this and our earlier work [14] also reveal distributions of small aggregates in our histograms (Fig. 1, Fig. S2), as recently also discussed by Kar et al. [37]; it would be of interest to see if a revised SP definition that lumps the concentrations of all the small aggregates could provide a more universal parameter to define phase transition thresholds.

The ideas revealed by our modeling and simulations can be tested experimentally, albeit at a more macroscopic scale. Experiments can be designed to titrate fluorescently labeled multivalent binding protein partners and use the fluorescence measured outside and inside condensates to determine the concentrations in each phase. Of course, our definition of SP is the product of monomer concentrations, while the dilute phase measured by fluorescence will contain a distribution of monomers and small oligomers. However, our studies suggest that the overall dilute phase concentration will scale with the monomer concentration so that the fluorescence of each monomer species will be proportional to monomer concentration. These experiments should be straightforward for in vitro validation of our predictions. The SP concept can then be marshalled as an additional tool to help elucidate the cellular formation of biomolecular condensates.

## Author contributions

A.C. designed the research, performed the simulations, built statistical tools, analyzed the data and wrote the paper. L.M.L. designed the research, supervised the project, secured the funding, analyzed the data, and wrote the paper.

## Acknowledgments

We gratefully acknowledge useful discussions with Michael Blinov. This work is supported by NIGMS grants R24 GM137787 and R01 GM132859.

## Video Legend

**Video 1: Clustering dynamics of ISD_2nm_rigid pair**. Total molecules = 160 + 160 = 320. Concentration ∼ 520 µM (last magenta point in Figure 8A).

