## Supplemental Figures for "The maximum solubility product marks the threshold for condensation of multivalent biomolecules"

### Supporting Material

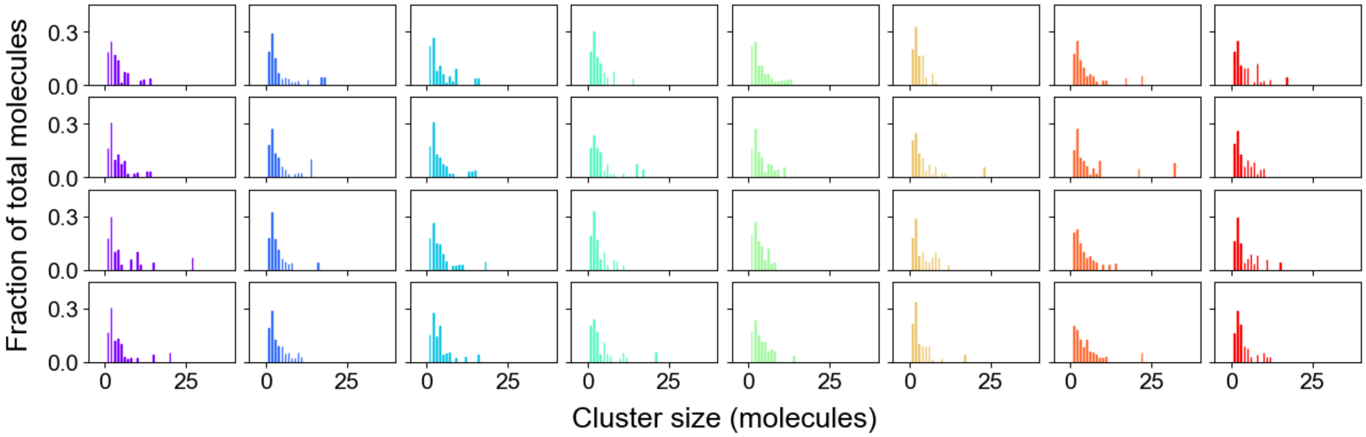


**Figure S1: Timecourse of cluster size distribution of a larger system below the phase transition threshold.** $A_5$ = $B_5$ = 100 µM (200 molecules). Each color (column) is a separate run and each row is a timepoint fluctuating around the steady state. 4 rows correspond to 10ms, 20ms, 30ms and 40ms respectively.

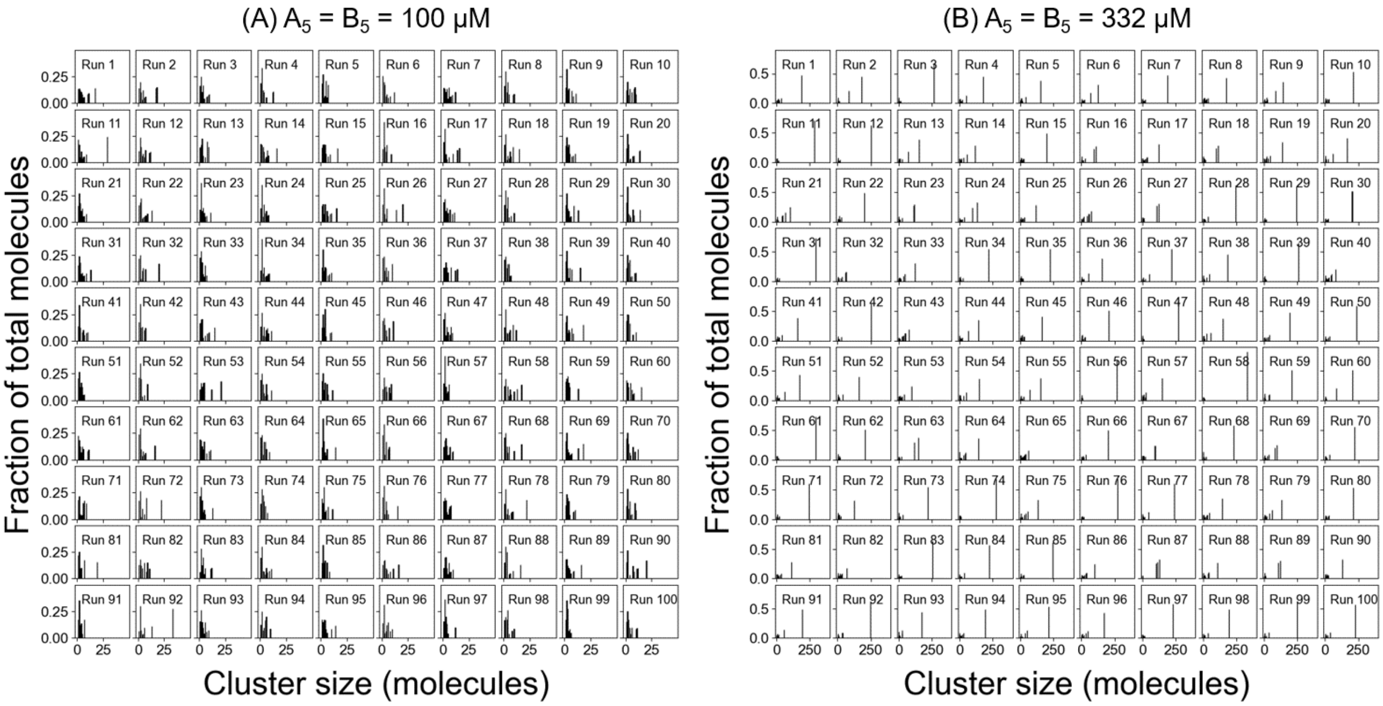


**Figure S2: Cluster size distributions (A) below and (B) above the phase transition threshold.** Only the last timepoint (40th ms) of 100 runs are displayed.

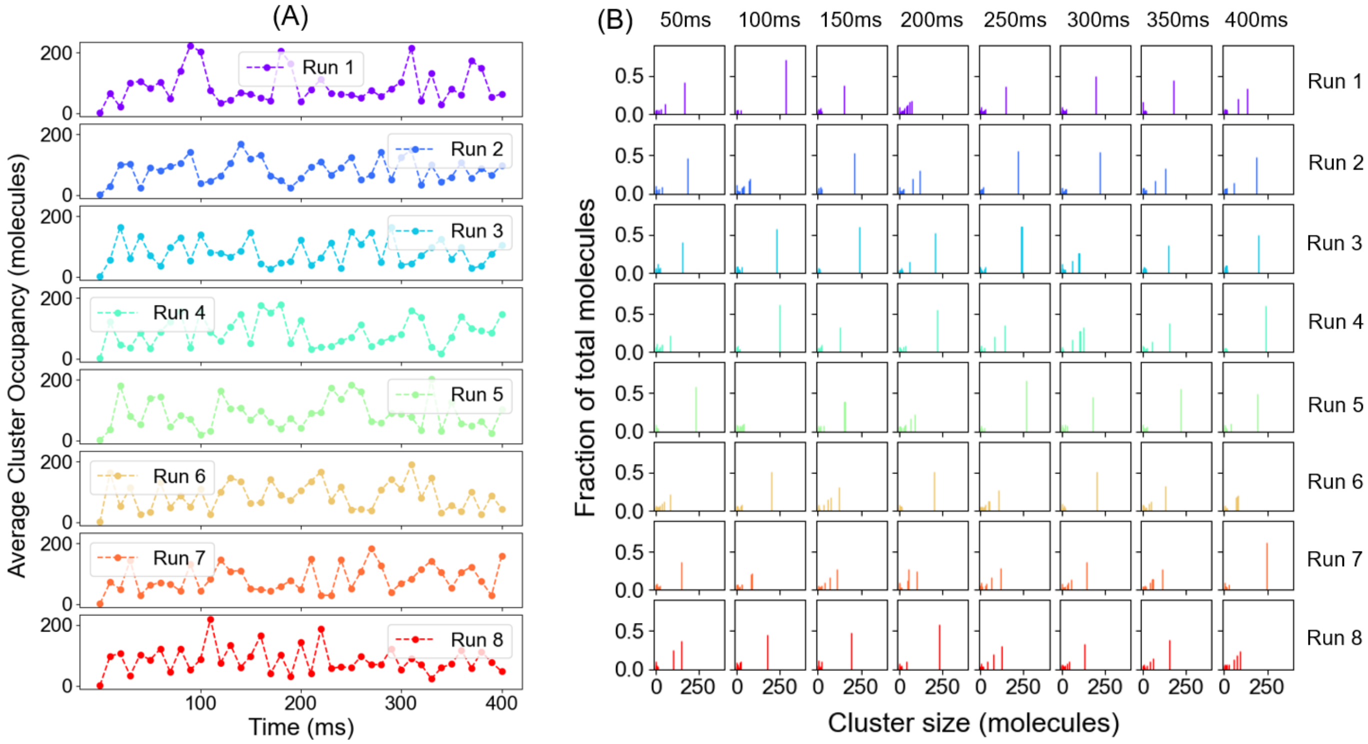


**Figure S3: Clustering dynamics shows similar behavior on longer time scale.** $A_5 = B_5 = 332\ \mu M$. Timecourse of (A) average cluster occupancy and (B) cluster distribution for 8 different runs.

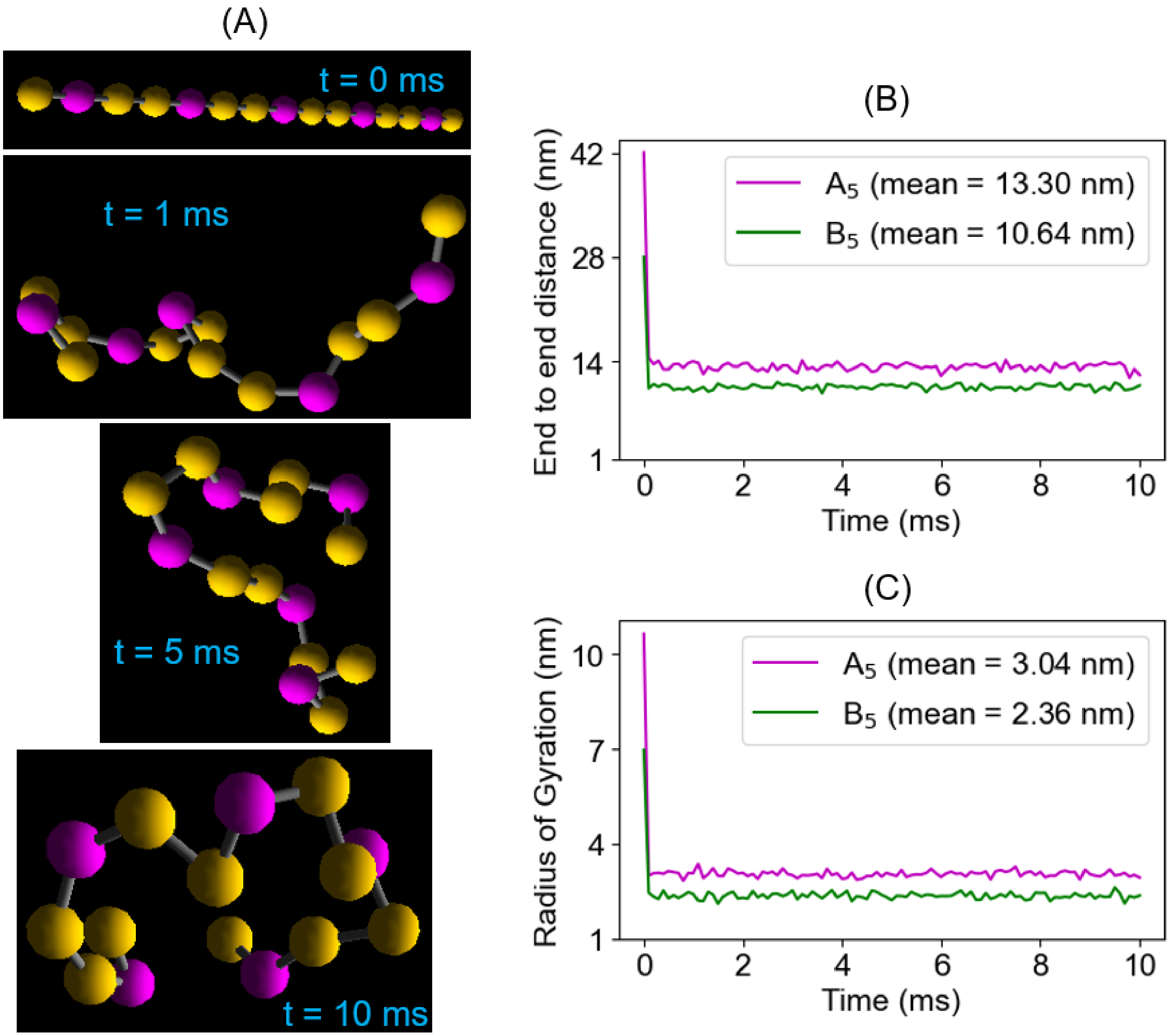


**Figure S4: Quantifying the compaction of flexible pentavalent monomers.** (A) Illustration of multiple conformations that a flexible molecule can take. The simulation starts with an elongated molecular structure and then relaxes to a compact state where the molecule samples multiple conformations around a steady degree of compaction. (B) Distance between the two terminal sites and (C) molecular radius of gyration, averaged over 100 trials. To quantify these parameters, we place a single molecule in a volume of 100*100*100 $nm^3$. Initial time point represents the linear molecular lengths ($A_5$ = 42nm and $B_5$ = 28nm). The mean values around the steady state are shown in the legends.

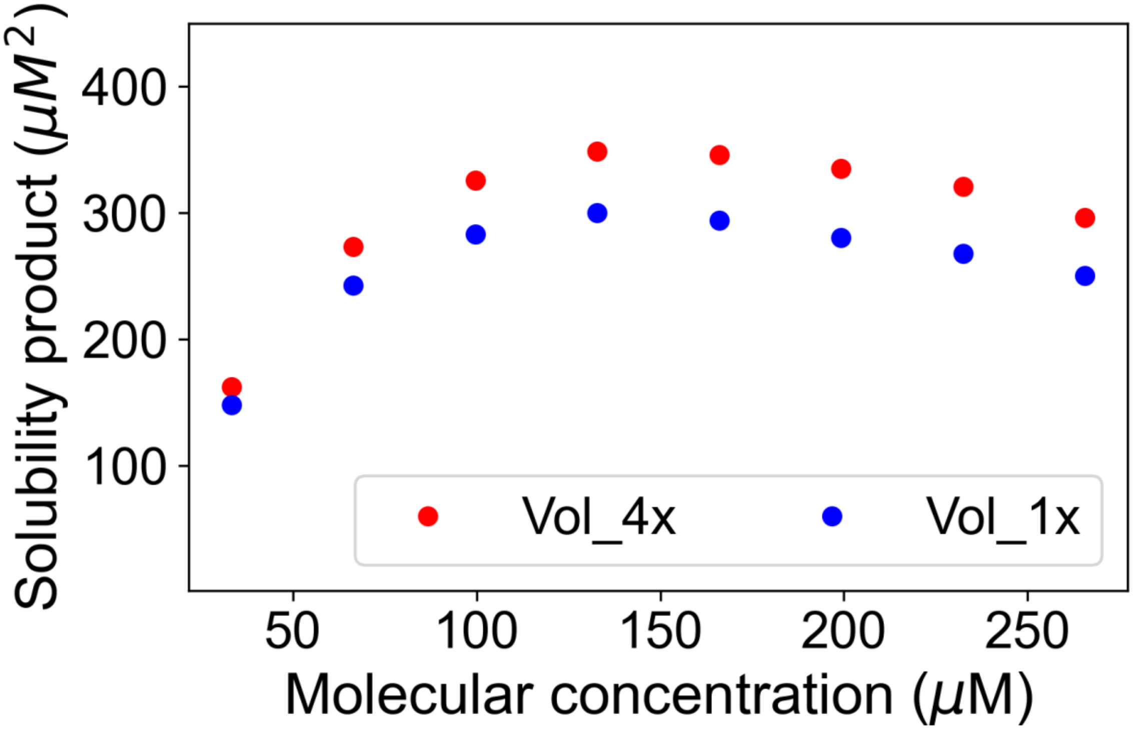


**Figure S5: Solubility product profile is qualitatively similar for a larger system.** Molecular pair described in Figure 1A is used here. Vol_1x = 100*100*100 $nm^3$ (Total molecules = 40, 80, 120, 160, 200, 240, 280, 320) and Vol_4x = 200*200*100 $nm^3$ (Total molecules = 160, 320, 480, 640, 800, 960, 1120, 1280).

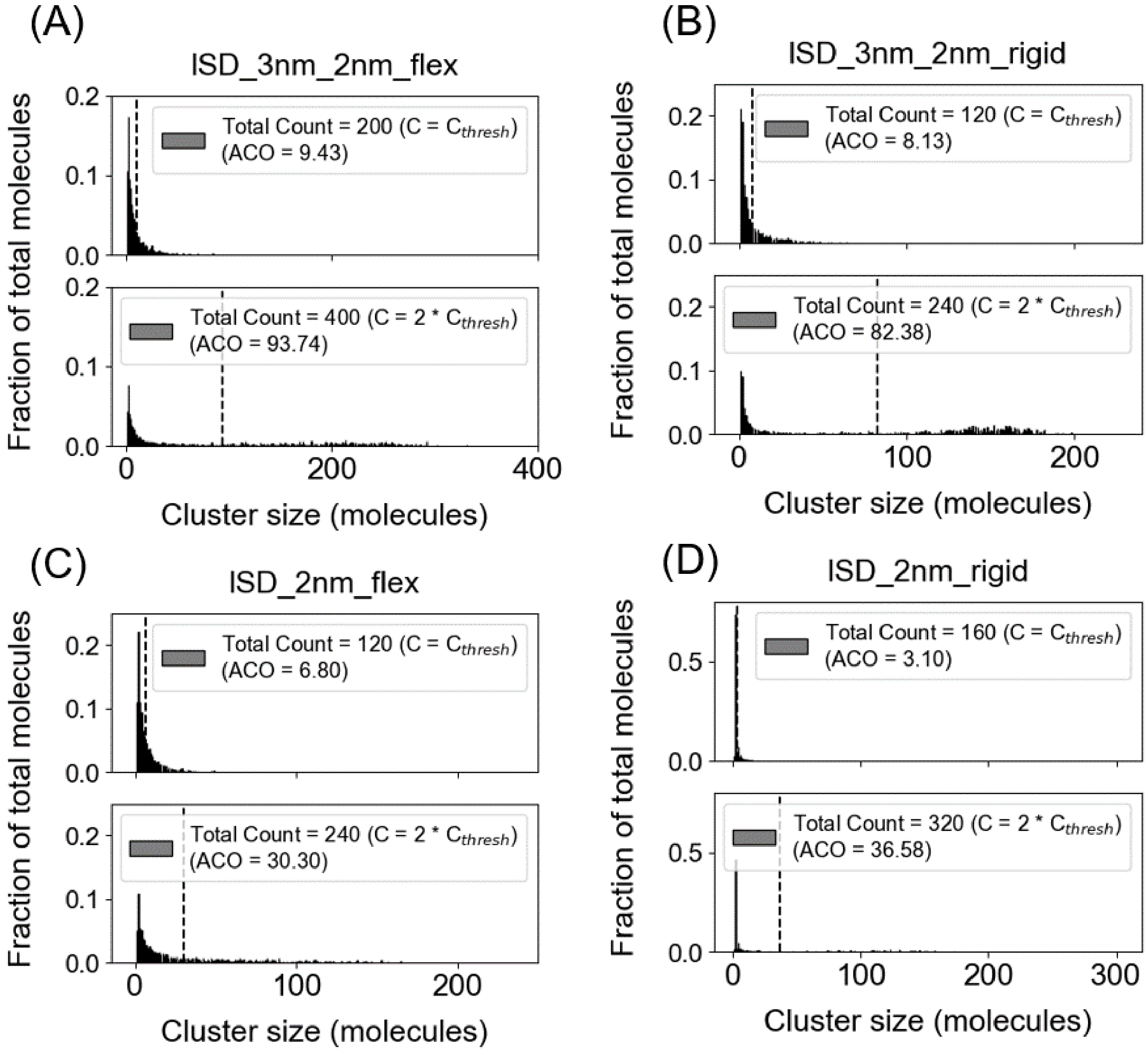


**Figure S6: Cluster size distributions for four structural pairs.** For each system, two concentration points are displayed.

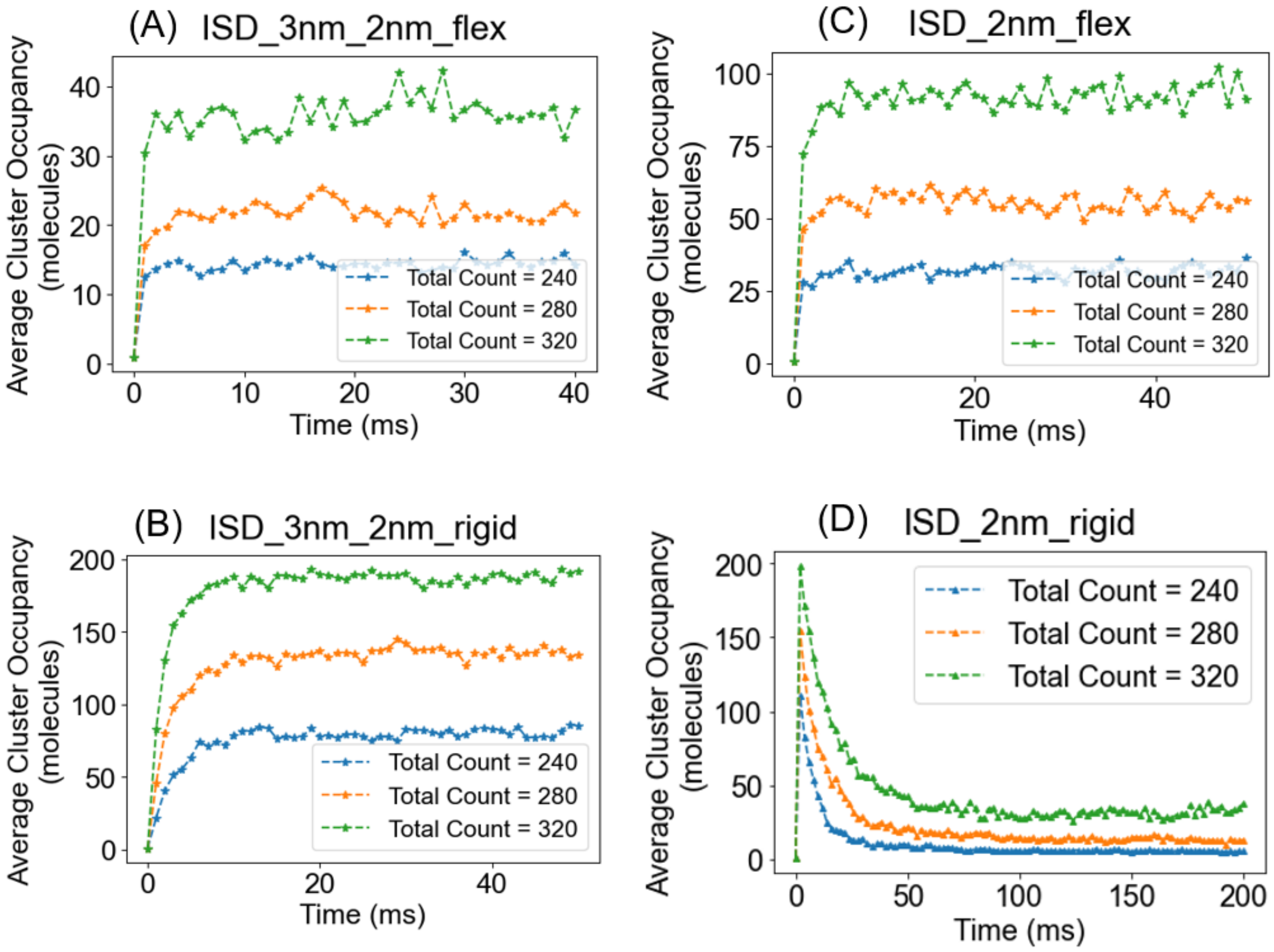


**Figure S7: Timecourse of average cluster occupancy for four structural pairs.** For each system, three different concentration plots are displayed. Relative molecular concentrations: (A) 1.5x, 1.75x, 2x (B) 2x, 2.33x, 2.66x (C) 2x, 2.33x, 2.66x (D) 1.2x, 1.4x, 1.6x where x is the threshold concentration of respective systems.

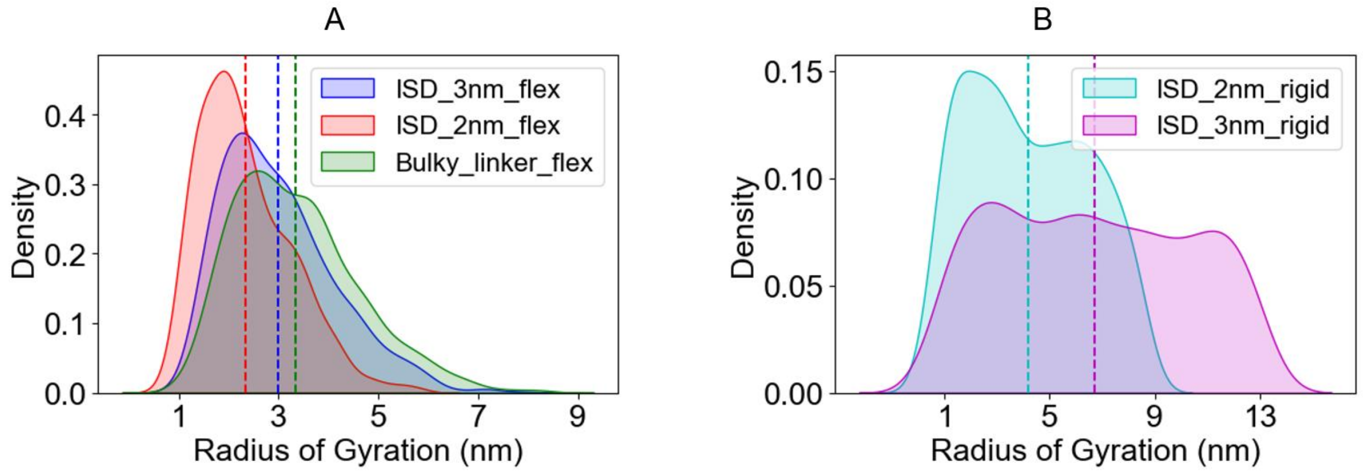


**Figure S8**: **Monomeric radius of gyration** for (A) three flexible molecules and (B) two rigid molecules. 1000 configurations are sampled over 100 stochastic trials. Dotted line indicates the mean of the distribution.

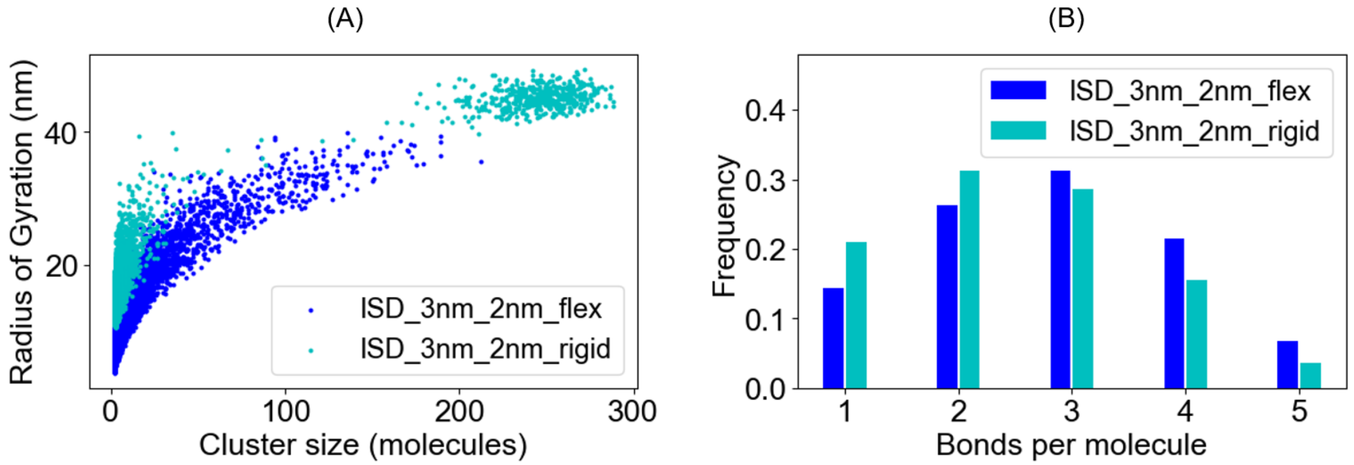


**Figure S9: Rigid monomers create extended clusters**. (A) Cluster specific radius of gyration and (B) Bond count distribution for flexible (Figure 1A) and corresponding rigid pair. Clusters are collected over 4 timepoints (10ms, 20ms, 30ms, 40ms) across 100 trials; total realizations = 400.
